# Generalizable sex differences in functional brain organization follow the cortical hierarchy in youth

**DOI:** 10.64898/2026.11.05.762913

**Authors:** Audrey C. Luo, Meike D. Hettwer, Aaron F. Alexander-Bloch, Joëlle Bagautdinova, Deanna M. Barch, Matthew Cieslak, Elizabeth A. Flook, Alexandre R. Franco, Raquel E. Gur, Ruben C. Gur, Arielle S. Keller, Gregory Kiar, Michael P. Milham, Laura Pritschet, Armin Raznahan, Taylor Salo, Giovanni A. Salum, Golia Shafiei, Russell T. Shinohara, Leah H. Somerville, Valerie J. Sydnor, Simon N. Vandekar, Xinyu Zhang, Sheila Shanmugan, Theodore D. Satterthwaite

**Author notes:** First Author: Audrey C. Luo. These authors contributed equally as senior authors (Sheila Shanmugan, Theodore D. Satterthwaite).

## Abstract

It remains unknown if sex differences in human functional connectivity follow a generalizable overarching principle. Here, we tested whether sex differences in functional connectivity during childhood and adolescence are organized along the sensorimotor-association (S-A) axis, a cortical hierarchy spanning unimodal sensorimotor to transmodal association cortex. We analyzed four large-scale, independent datasets of youth ages 5-23 years (total *N* = 3,355). Across datasets, males showed greater connectivity between functional networks, whereas females showed greater connectivity within networks, most prominently within the default mode network. More broadly, the sensorimotor and association ends of the S-A axis were more segregated in females and more integrated in males. This difference extended to macroscale cortical organization: the cortical hierarchy was expanded in females relative to males along the principal gradient of functional connectivity. Together, these findings identify the S-A axis as a generalizable organizing principle of sex differences in functional brain organization during youth.

## INTRODUCTION

Sex is a major source of individual variation in the developing human brain, and understanding how brain organization differs by sex is fundamental to characterizing typical development. The clinical relevance of understanding sex differences is underscored by the marked sex differences in many psychiatric disorders (*1*). Human neuroimaging studies have identified sex differences in multiple features of brain structure and function, including large-scale patterns of resting-state functional connectivity across cortical regions, functional networks, and individualized network topography (*2–7*). However, these differences have been characterized using disparate measures of functional organization. This heterogeneity makes findings difficult to integrate across studies and obscures whether sex differences in functional connectivity follow an overarching organizing principle across the cortex. Moreover, most studies have relied on single datasets, limiting our understanding of whether these findings generalize across independent datasets (*8*, *9*), and few studies have evaluated their consistency across heterogeneous samples (*6*, *10*). Here, we asked whether sex differences in functional connectivity in youth are systematically organized along a cortical hierarchy.

Several findings suggest that sex differences in functional connectivity are not randomly distributed but rather exhibit consistent patterning across the cortex. In a large sample of youth, males showed greater connectivity between functional networks, whereas females showed greater connectivity within networks (*7*), a pattern also observed in adults (*11*). Related studies across the lifespan have reported greater functional network segregation in females, particularly in association networks (*12*, *13*). Studies using personalized functional networks have also identified prominent sex differences in the topography of association networks (*5*, *10*). Furthermore, sex differences in the default mode network have been repeatedly observed in both functional network topography and connectivity, with several studies reporting greater within-network connectivity in females (*3*, *5*, *10*, *14*, *15*). However, the magnitude and direction of sex differences in default mode network organization have varied across studies. Sex differences have also been reported in control, attention, sensorimotor, visual, and limbic systems (*3*, *5*, *10*, *16*), but it is unclear how these findings relate to one another. At present, the field lacks a broader organizing framework that parsimoniously unifies disparate existing results.

The sensorimotor-association (S-A) axis provides a potential framework for integrating these findings (*17*). This major axis of cortical organization reflects a cortical hierarchy spanning regions involved in unimodal sensory and motor processing to higher-order association regions supporting complex cognition and socioemotional processing (*18*). The S-A axis captures broad variation in cortical anatomy and function by integrating ten multimodal cortical maps, including the principal gradient of functional connectivity (*17*, *19*). Developmental changes in functional connectivity also follow this hierarchy, with maturational patterns varying systematically from sensorimotor to association cortex (*13*, *20*). Recent work in young adults has identified sex differences in functional organization along the S-A axis (*21*). Together, these findings motivate testing whether sex differences in functional connectivity follow the S-A axis and generalize across independent datasets of youth.

To address this question, we analyzed four independent, large-scale datasets (total *N* = 3,355) of youth ages 5-23 years from diverse populations and geographic regions of the United States. We hypothesized that sex differences in functional connectivity would be organized along the S-A axis. To evaluate sex differences across complementary levels of functional organization, we examined functional connectivity at the regional, network, and edge levels, as well as macroscale organization along the principal gradient of functional connectivity. At each level, we evaluated whether these effects generalized across all four datasets. As described below, results reveal generalizable sex differences in functional connectivity that align with the S-A axis. These findings extend to macroscale cortical organization: the cortical hierarchy was expanded in females relative to males along the principal gradient.

## RESULTS

To characterize sex differences in functional connectivity in youth, we used resting-state and task-based functional MRI from four independent datasets (total *N* = 3,355; 1,608 female): the Philadelphia Neurodevelopmental Cohort (PNC; *n* = 1,207; 646 female; ages 8-23), the Nathan Kline Institute-Rockland Sample (NKI; *n* = 397; 186 female; ages 6-22), the Human Connectome Project: Development (HCP-D; *n* = 625; 337 female; ages 5-22), and the Healthy Brain Network (HBN; *n* = 1,126; 439 female; ages 5-22) (see Table 1 of (*20*)). These datasets were previously used to characterize the development of functional connectivity along the sensorimotor-association (S-A) axis (*20*). We additionally analyzed a pooled dataset that combined participants from all four datasets. Measures were harmonized across sites in the multi-site datasets (HCP-D and HBN) and across both sites and datasets in the pooled dataset. We examined sex differences in functional connectivity at the region, network, and edge levels, as well as in the macroscale organization of cortical connectivity captured by the principal functional gradient. For each measure, we fit generalized additive models (GAMs) with sex, age as a smooth term, and in-scanner head motion as a covariate. Sex effects were quantified using the robust effect size index (RESI), which is comparable across model types and independent of sample size (*22*, *23*); negative values indicate higher values in males and positive values indicate higher values in females. According to (*23*), absolute RESI values of 0-0.1, 0.1-0.25, 0.25-0.4, and > 0.4 can be interpreted as no-to-small, small-to-medium, medium-to-large, and large effects, respectively. Our cortical parcellation was the Schaefer 200 atlas with a 17-network partition based on the 17-network Yeo solution (*24*, *25*).

At the region level, we first examined whether sex differences in functional connectivity (FC) strength replicated across datasets. We then decomposed each region’s FC strength into between- and within-network connectivity. We contextualized the resulting sex differences using Neurosynth meta-analytic maps of cognitive terms. At the network level, we evaluated sex differences in connectivity for every pair of the 17 functional networks, including each network’s connectivity with itself. At the edge level, we tested whether sex differences in connectivity between pairs of regions were organized along the S-A axis. Finally, given the edge-level findings, we examined whether the cortical hierarchy as a whole was compressed in males and expanded in females, as measured by the distance between networks along the principal gradient of functional connectivity.

### Replicable sex differences in FC strength reflect greater between-network connectivity in males and greater within-network connectivity in females

We first sought to characterize sex differences in FC strength, defined as the mean edge strength between a given region and all other cortical regions, and to evaluate how consistent these differences were across datasets. In every dataset, males exhibited greater FC strength than females across widespread cortical regions (**Fig. 1A**). Significant sex differences in FC strength (FDR-corrected, *Q*_FDR_ < 0.05) were present in 154 of 200 regions (77%) in the PNC, 33 (17%) in NKI, 56 (28%) in HCP-D, 63 (32%) in HBN, and 178 (89%) in the pooled dataset. With the exception of a single region in one dataset, all significant effects reflected greater FC strength in males. We note that fewer regions reached significance in NKI, almost certainly due to the reduced statistical power of this smaller sample. The spatial patterns of FC strength sex effects were significantly correlated across datasets (*r* = 0.29 to 0.56, all *p*_spin_ < 0.001; **Fig. S1A**). As a descriptive summary, we tallied the number of datasets in which each region showed a significant sex difference (**Fig. 1B**). Eighty-five regions (43%) showed significantly greater FC strength in males in two or more datasets and 39 (20%) in three or more. Fourteen regions were significant in all four datasets, including posteromedial (precuneus, posterior cingulate, and retrosplenial) and prefrontal cortices. All 14 regions were among the 20% with the largest mean effect sizes across datasets.

**Figure 1.**
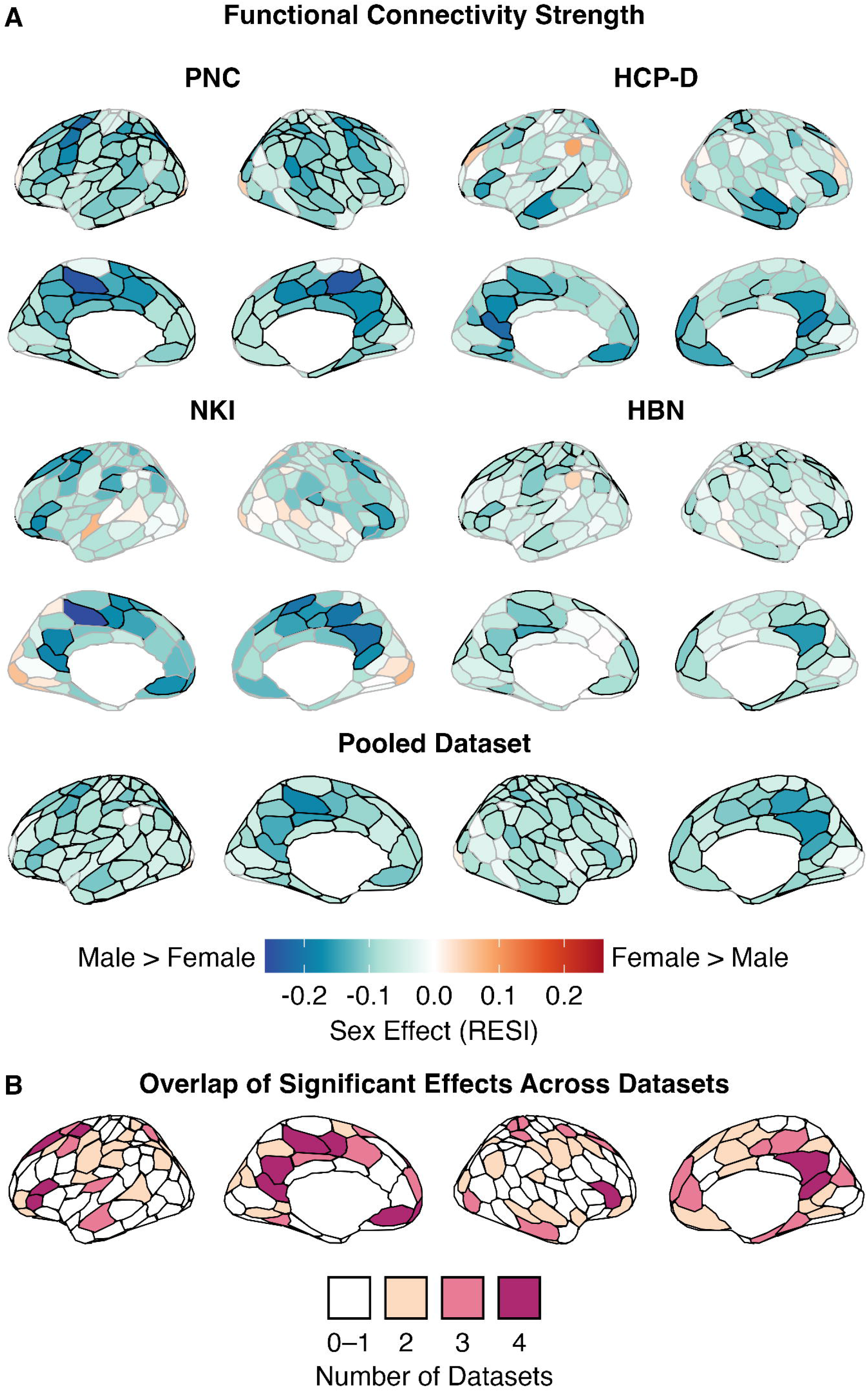
Males exhibit greater functional connectivity strength across the cortex. **(A)** Sex effects on functional connectivity (FC) strength, quantified using the robust effect size index (RESI), are shown on the cortical surface for the PNC, NKI, HCP-D, and HBN. In every dataset, males exhibit greater FC strength across widespread cortical regions. A pooled analysis combining participants from all four datasets confirms this effect across nearly the entire cortex. Blue (negative RESI) indicates greater connectivity in males, whereas red (positive RESI) indicates greater connectivity in females. Regions outlined in black display significant sex differences in FC strength (*Q*_FDR_ < 0.05). **(B)** To assess the consistency of sex effects across datasets, cortical surfaces show the number of datasets among PNC, NKI, HCP-D, and HBN in which each region displayed a significant sex difference in FC strength. Regions with significantly greater FC strength in males in two or more datasets span much of the cortex, with those significant in all four datasets concentrated in the posteromedial and prefrontal cortices. No region exhibits significantly greater FC strength in females than males in more than one dataset. Results are shown in the Schaefer 200 atlas.

While FC strength summarizes the average connectivity of a given region to the rest of the cortex, it does not capture whether sex differences in these connections fall within or outside of that region’s functional network. To determine which connections underlie greater FC strength in males, we separated FC strength into its two constituent components: between-network and within-network connectivity, defined as the mean edge strength of a given region to all other regions outside of and within that region’s functional network, respectively.

Sex differences in between-network connectivity closely resembled those in FC strength (**Fig. 2A**). Significant effects were present in 149 regions (75%) in the PNC, 28 (14%) in NKI, 73 (37%) in HCP-D, 78 (39%) in HBN, and 176 (88%) in the pooled dataset. Nearly all significant effects showed greater connectivity in males than females. No region showed significantly greater between-network connectivity in females in more than one dataset. Sex effects on between-network connectivity were spatially correlated across datasets (*r* = 0.42 to 0.61; all *p*_spin_ < 0.001; **Fig. S1B**). Ninety-eight regions (49%) showed significantly greater between-network connectivity in males in two or more datasets and 52 (26%) in three or more. The 16 regions significant in all four datasets were concentrated in posteromedial and prefrontal cortices and included all 14 regions showing replicable sex differences in FC strength (**Fig. 2B**); all 16 regions also ranked within the top 20% of mean effect sizes across datasets. To contextualize these effects, we focused on regions showing significant sex differences in two or more datasets and averaged their sex effects across datasets (**Fig. S2A-B**). We then used Spearman rank correlations to compare the spatial pattern of sex effects with Neurosynth meta-analytic maps of 125 cognitive terms (*26*, *27*). Significance was assessed with spin-based spatial null models and FDR correction across terms (*28*). These correlations index spatial correspondence with task-activation maps and do not establish sex differences in the corresponding cognitive functions, which were not directly evaluated in the present analyses. Further, these correlations do not imply that greater connectivity in either sex reflects better or worse performance in these domains. Regions with greater between-network connectivity in males aligned with four terms: cognitive control, thought, utility, and task difficulty (*r* = −0.43 to −0.33; all *Q*_FDR_ < 0.05; **Fig. S2C**).

**Figure 2.**
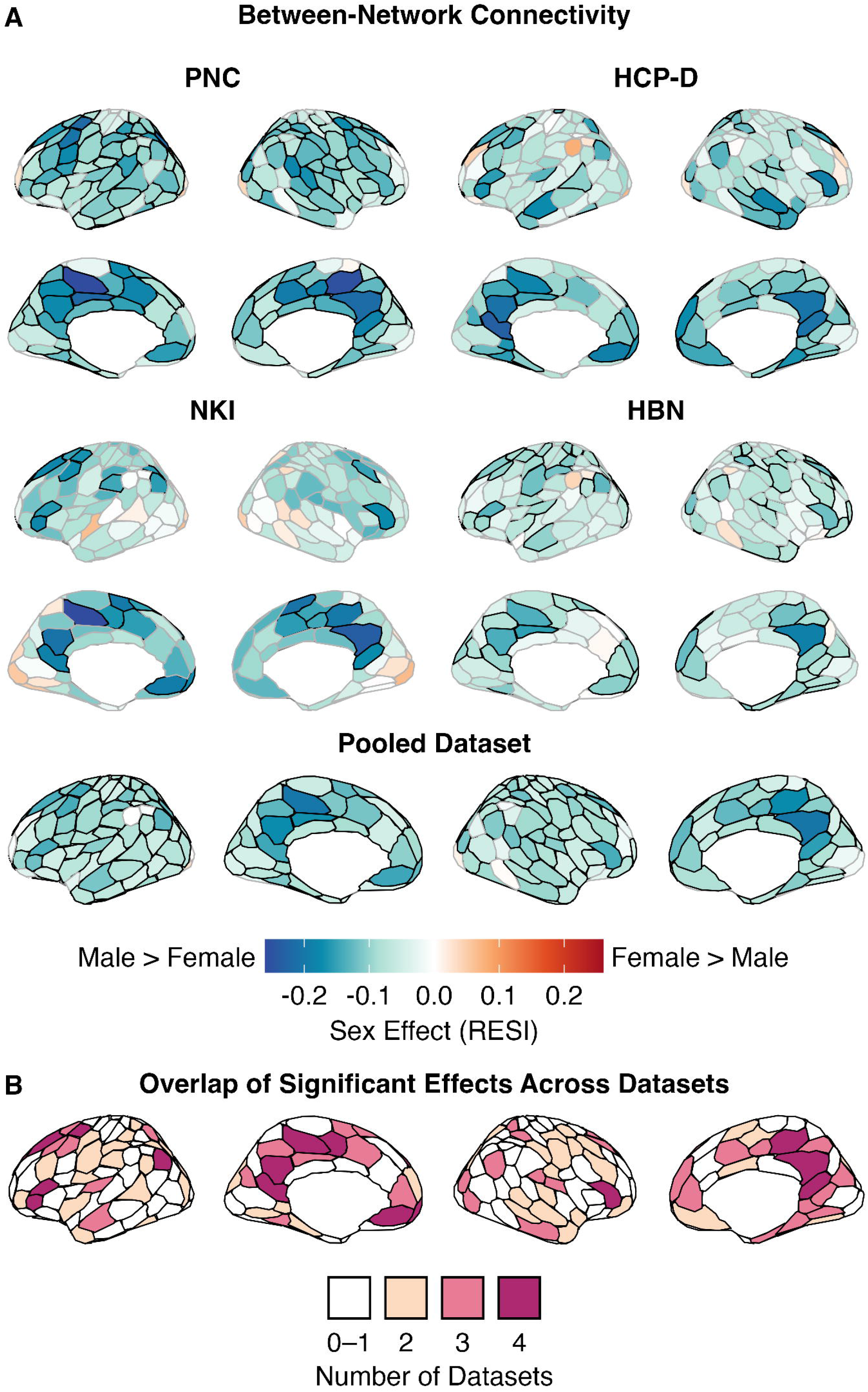
Greater between-network connectivity contributes to greater functional connectivity strength in males. **(A)** Sex effects (RESI) on between-network connectivity are shown for PNC, NKI, HCP-D, and HBN (Schaefer 200). In every dataset, males exhibit greater between-network connectivity across widespread cortical regions. A pooled analysis reveals greater between-network connectivity in males across nearly the entire cortex. Regions with significant sex differences (*Q*_FDR_ < 0.05) are outlined in black. **(B)** Cortical surfaces display the number of datasets in which each region showed a significant sex difference in between-network connectivity. Regions with significantly greater between-network connectivity in males (negative RESI) in two or more datasets are distributed throughout the cortex, whereas those significant in all four datasets are concentrated in the posteromedial and prefrontal cortices. No region shows significantly greater between-network connectivity in females (positive RESI) in more than one dataset.

In contrast to the distributed greater between-network connectivity in males, sex differences in within-network connectivity predominantly reflected greater connectivity in females (**Fig. 3A**). Significant effects were present in 73 regions (37%) in the PNC, 12 (6%) in NKI, 45 (23%) in HCP-D, 35 (18%) in HBN, and 102 (51%) in the pooled dataset. Sex effects on within-network connectivity were highly spatially correlated across datasets (*r* = 0.68 to 0.82; all *p*_spin_ < 0.001; **Fig. S1C**). In total, 32 (16%) regions showed significantly greater within-network connectivity in females in two or more datasets and 16 (8%) in three or more, concentrated in the medial prefrontal, posterior cingulate, precuneus, and inferior parietal cortices (**Fig. 3B**). The six regions significant in all four datasets ranked within the top 20% of mean effect sizes across datasets. Regions with greater within-network connectivity in females in two or more datasets aligned with Neurosynth maps for nine cognitive terms, including autobiographical memory, social cognition, belief, and intention (*r* = 0.42 to 0.58; all *Q*_FDR_ < 0.05; **Fig. S2D**). Conversely, 15 regions showed significantly greater within-network connectivity in males in two or more datasets, primarily in the temporal pole and somatomotor cortex.

**Figure 3.**
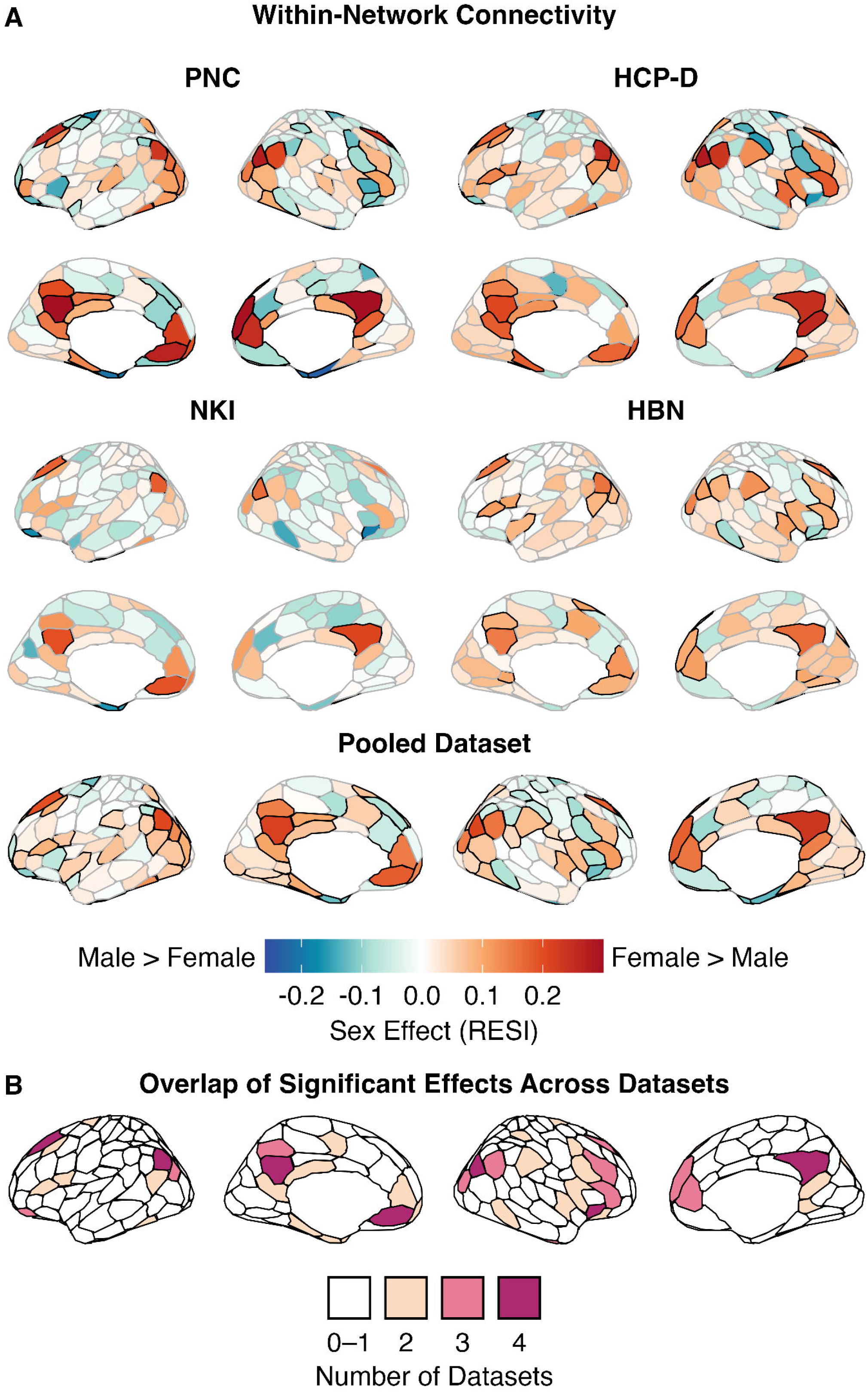
Females exhibit greater within-network connectivity in medial prefrontal, posterior cingulate, precuneus, and inferior parietal cortices. **(A)** Sex effects (RESI) on within-network connectivity are displayed for the PNC, NKI, HCP-D, and HBN (Schaefer 200). Females show greater within-network connectivity than males in most regions, with the largest effects in the medial prefrontal, posterior cingulate, precuneus, and inferior parietal cortices. This spatial pattern is also evident in the pooled dataset. Regions outlined in black show significant sex differences (*Q*_FDR_ < 0.05). **(B)** The number of datasets in which each region showed a significant sex difference is displayed on the cortical surface. Regions with significantly greater within-network connectivity in females (positive RESI) in two or more datasets are located predominantly in the medial prefrontal, posteromedial (precuneus, posterior cingulate, and retrosplenial), and inferior parietal cortices. Regions with significantly greater within-network connectivity in males (negative RESI) in two or more datasets are located in the temporal pole and in somatomotor, insular, prefrontal, orbitofrontal, and attention-related cortices.

Having established that sex has a main effect on functional connectivity, we next evaluated if sex differences varied with age. For this question, we focused on the pooled dataset to maximize statistical power. Even in the large pooled dataset, we found that sex differences in FC strength and between-network connectivity did not vary with age. However, age-by-sex interactions in within-network connectivity were observed in 22 regions (11%), including default mode subnetwork A regions, where connectivity increased more with age in females than in males; in somatomotor, salience/ventral attention, and limbic temporal pole regions, connectivity increased more with age in males (**Fig. S3**). Interactions in individual datasets were sparse and did not replicate.

Taken together, these results show that greater FC strength in males replicates across four independent datasets and reflects opposing effects in between- and within-network connectivity. Males showed greater between-network connectivity across widespread cortical regions, whereas greater within-network connectivity in females was largely confined to default mode regions, most of which also showed greater between-network connectivity in males.

### Network-level sex differences converge on default mode subnetwork A

To identify the functional networks to which the region-level effects belonged, we next examined sex differences in connectivity for all 153 pairs of the 17 functional networks, including each network with itself (**Fig. 4A**). Network-pair connectivity was computed by averaging connectivity across all pairs of regions belonging to the two networks, or across all pairs of regions within the same network. Significant effects were present in 76 of 153 network pairs (50%) in the PNC, 6 (4%) in NKI, 43 (28%) in HCP-D, 31 (20%) in HBN, and 95 (62%) in the pooled dataset. Males showed greater connectivity in most of the significant between-network pairs, consistent with the widespread greater between-network connectivity observed at the region level. Connectivity within default mode subnetwork A – which includes medial and dorsal prefrontal, posterior cingulate/precuneus, and inferior parietal cortices – was greater in females and was the largest effect in this direction in the PNC (RESI = 0.33, *Q*_FDR_ < 0.0001), NKI (RESI = 0.18, *Q*_FDR_ = 0.014), HBN (RESI = 0.16, *Q*_FDR_ < 0.0001), and the pooled dataset (RESI = 0.23, *Q*_FDR_ < 0.0001). In HCP-D, it was the second largest (RESI = 0.24, *Q*_FDR_ < 0.0001), after connectivity within default mode subnetwork C (RESI = 0.26, *Q*_FDR_ < 0.0001). Connectivity within default mode subnetwork A was also one of 10 network pairs with a significant age-by-sex interaction in the pooled dataset, where connectivity increased more with age in females than in males (**Fig. S4**).

**Figure 4.**
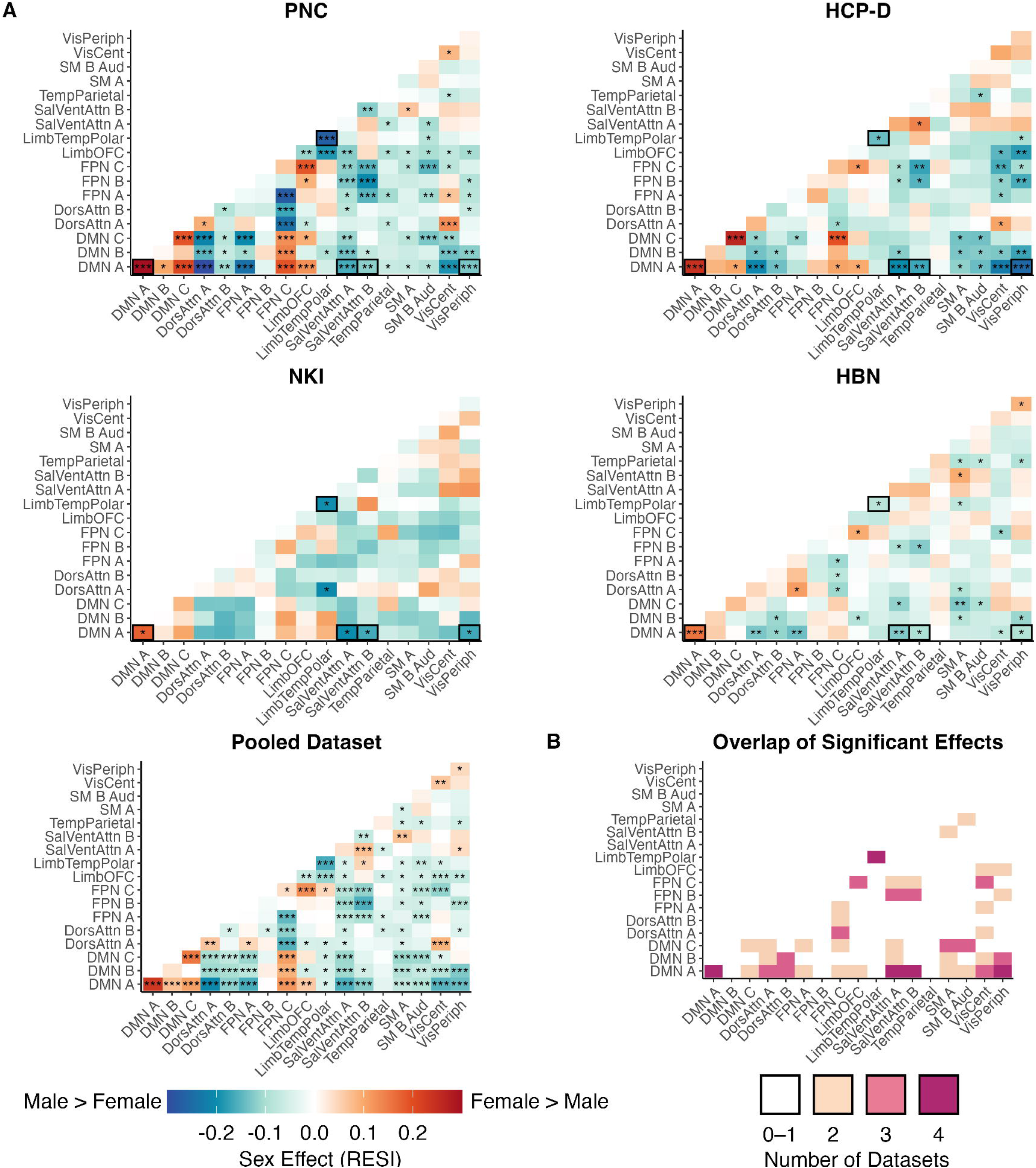
Sex differences in default mode subnetwork A connectivity replicate across datasets. (**A**) Sex effects (RESI) on connectivity between each pair of functional networks (Yeo 17-network partition; 153 network pairs) are shown for the PNC, NKI, HCP-D, HBN, and the pooled dataset. Connectivity was averaged across all pairs of regions belonging to two different networks (off-diagonal cells) or to the same network (diagonal cells). Connectivity within default mode subnetwork A is greater in females and is the largest female-greater effect in the PNC, NKI, HBN, and the pooled dataset, and the second largest in HCP-D (after default mode subnetwork C). Connectivity between default mode subnetwork A and attention or visual networks is greater in males in most datasets. Asterisks denote significance after FDR correction (\**Q*_FDR_ < 0.05, \*\**Q*_FDR_ < 0.001, \*\*\**Q*_FDR_ < 0.0001). In individual dataset panels, black outlines mark network pairs significant in all four datasets. **(B)** The matrix displays the number of datasets in which each network pair showed a significant sex difference in either direction (*Q*_FDR_ < 0.05). Five network pairs show significant sex differences in all four datasets. Connectivity within default mode subnetwork A is greater in females. Males show greater connectivity within the limbic temporal pole network and between default mode subnetwork A and the salience/ventral attention A, salience/ventral attention B, and peripheral visual networks. DMN, default mode network; DorsAttn, dorsal attention; FPN, frontoparietal control; LimbOFC, limbic orbitofrontal; LimbTempPolar, limbic temporal pole; SalVentAttn, salience/ventral attention; TempParietal, temporal parietal; SM A, somatomotor A; SM B Aud, somatomotor B/auditory; VisCent, central visual; VisPeriph, peripheral visual.

To assess the consistency of these effects, we counted the number of datasets in which each network pair showed a significant sex difference (**Fig. 4B**). Five network pairs showed significant sex differences in all four datasets. In addition to greater connectivity within default mode subnetwork A in females, males showed greater connectivity between default mode subnetwork A and three other networks in every dataset (all *Q*_FDR_ < 0.05): salience/ventral attention A (RESI = −0.20 to −0.13), salience/ventral attention B (RESI = −0.17 to −0.09), and peripheral visual (RESI = −0.25 to −0.09) networks. Males also showed greater connectivity within the limbic temporal pole network (RESI = −0.27 to −0.08; all *Q*_FDR_ < 0.05). All five pairs ranked within the top 20% of mean effect sizes across datasets. These findings identify default mode subnetwork A as a point of convergence for sex differences: it was involved in four of the five significant network pairs that replicated across all four datasets. Located near the association pole of the cortical hierarchy, default mode subnetwork A showed greater connectivity within the network in females and greater connectivity with salience/ventral attention and visual networks in males.

### Sex differences in edge-level connectivity are organized along the sensorimotor-association axis

Given that the most reproducible network-level sex differences converged on default mode subnetwork A, a higher-order association network, we next asked whether sex differences in connectivity follow a more general hierarchical pattern across the cortex. To test this, we annotated each functional connection (edge) with the sensorimotor-association (S-A) axis ranks (*17*) of the two regions it connects. We modeled the effect of each combination of S-A axis ranks, representing pairs of brain regions, on edge-level sex effects using a tensor product smooth (**Fig. 5**). This allowed us to examine how sex differences vary with the hierarchical span of connections and the positions of connected regions along the S-A axis.

**Figure 5.**
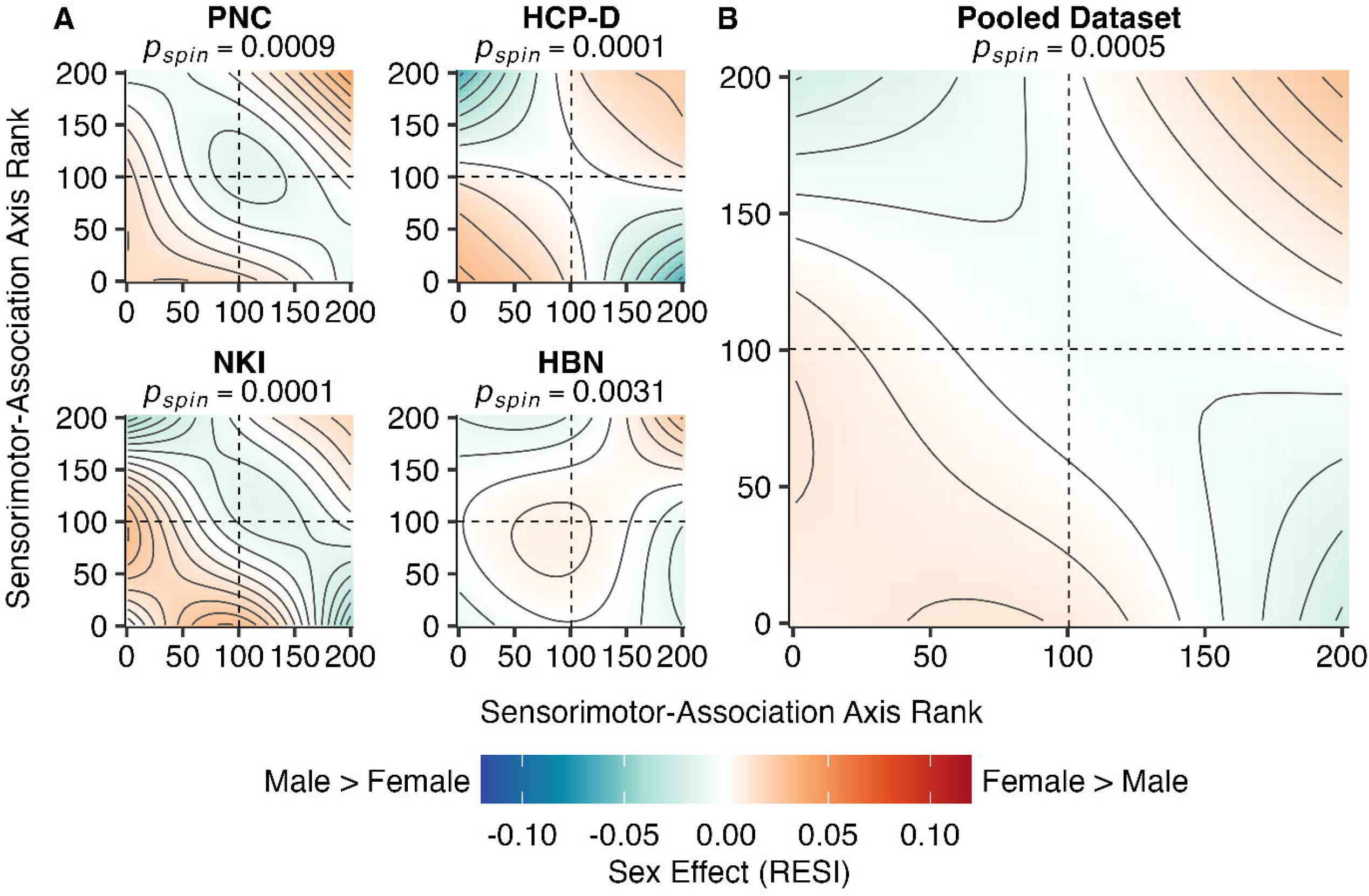
Sex differences in edge-level connectivity are organized along the sensorimotor-association axis. **(A and B)** Topographical plots display edge-level sex effects (RESI) across the sensorimotor-association (S-A) axis for PNC, NKI, HCP-D, and HBN **(A)** and the pooled dataset **(B)**. Low ranks correspond to sensorimotor regions and high ranks to association regions. Colors show sex effects fitted with a tensor product smooth of the S-A axis ranks of each edge’s connected regions. Contour lines connect points with equal sex effects. Females exhibit greater connectivity between regions at similar positions on the axis (sensorimotor-to-sensorimotor, bottom left of each panel; association-to-association, top right). Males show greater connectivity between regions at opposite ends of the axis (top left and bottom right of each panel). In every dataset, the S-A axis ranking of regions connected by each edge significantly explains variance in edge-level sex effects (PNC: *p*_spin_ = 0.0009; NKI: *p*_spin_ = 0.0001; HCP-D: *p*_spin_ = 0.0001; HBN: *p*_spin_ = 0.0031; pooled dataset: *p*_spin_ = 0.0005). Spin tests assessed the statistical significance of the variance explained by the tensor product smooth.

Across datasets, females showed greater edge-level connectivity between regions with similar S-A ranks, specifically in sensorimotor-to-sensorimotor and association-to-association connections. In contrast, males showed greater connectivity between regions at opposite ends of the S-A axis (**Fig. 5A**). This pattern was also evident in the pooled dataset (**Fig. 5B**). The S-A axis ranks of connected regions explained significant variance in edge-level sex effects in every dataset, as assessed by spin-based spatial null models (PNC: *p*_spin_ = 0.0009; NKI: *p*_spin_ = 0.0001; HCP-D: *p*_spin_ = 0.0001; HBN: *p*_spin_ = 0.0031; pooled dataset: *p*_spin_ = 0.0005). Together, these results reveal a broader hierarchical pattern underlying sex differences in functional connectivity: the sensorimotor and association ends of the axis were more segregated from each other in females and more integrated in males.

### Females exhibit a more expanded cortical hierarchy than males

Given that edge-level connectivity was stronger within each end of the S-A axis in females and between its opposite ends in males, connectivity profiles at the sensorimotor and association ends of this axis may be more distinct from one another in females than in males. We therefore asked whether higher-order regions were more differentiated from lower-order regions in females, corresponding to a more expanded cortical hierarchy in females and a more compressed hierarchy in males. To this end, we computed the principal gradient of functional connectivity for each participant using principal component analysis, aligned to a group template (*19*). This gradient orders regions according to the similarity of their connectivity profiles and recapitulates the S-A axis, spanning from sensorimotor to association cortex in every dataset and in both sexes (**Fig. S5**). Each region’s loading thus indicates its position along the hierarchical principal gradient. We first quantified global dispersion as the overall spread of regions along the principal gradient, such that greater global dispersion indicates a more expanded hierarchy. To examine how networks were positioned relative to one another on this gradient, we defined each network’s position as its centroid, the median loading of its regions. We quantified between-network dispersion as the distance between network centroids, or the absolute difference between median loadings of each network, along the principal gradient (*21*, *29*). Greater values indicate greater differentiation between networks in their respective principal gradient positions.

Global dispersion was greater in females in the PNC (RESI = 0.09, *p* = 0.0018), NKI (RESI = 0.12, *p* = 0.019), HCP-D (RESI = 0.19, *p* < 0.0001), and the pooled dataset (RESI = 0.10, *p* < 0.0001), but not significantly in HBN (RESI = 0.05, *p* = 0.1; **Fig. S6**), consistent with greater overall expansion of the cortical hierarchy in females. Significant sex differences in between-network dispersion were present in 59 of 136 network pairs (43%) in the PNC, 22 (16%) in NKI, 65 (48%) in HCP-D, 8 (6%) in HBN, and 89 (65%) in the pooled dataset (**Fig. 6A**). Dispersion was greater in females for most significant network pairs (>70%) in every dataset. Significantly greater dispersion in males was rare (0-4 pairs per individual dataset) and did not replicate across datasets. The largest effects were predominantly between higher-order association networks (e.g., default mode and frontoparietal control subnetworks) and lower-order networks (largest effect per dataset: RESI = 0.12 to 0.23, all *Q*_FDR_ < 0.05). Greater dispersion in females was also consistent across datasets: 24 network pairs showed significant effects in three or more datasets (**Fig. 6B**). Most of these pairs consisted of a higher-order default mode, frontoparietal control, or limbic orbitofrontal network and lower-order somatomotor, auditory, visual, or salience/ventral attention network. Among these, dispersion between default mode subnetwork A and salience/ventral attention A was greater in females in every dataset except HBN (RESI = 0.14 to 0.21, all *Q*_FDR_ < 0.05), as was dispersion between default mode subnetworks A and C and the lowest-order networks (somatomotor B/auditory, central visual, and peripheral visual; RESI = 0.11 to 0.23, all *Q*_FDR_ < 0.05). For default mode subnetwork A, these same network pairs showed greater connectivity in males in most or all datasets (see **Fig. 4**, above). Thus, default mode subnetwork A was more differentiated from lower-order networks in females, both in its connectivity and in its position along the principal gradient. Together, these results indicate that the cortical hierarchy is expanded in females, with greater differentiation between higher-order association and lower-order networks along the principal gradient.

**Figure 6.**
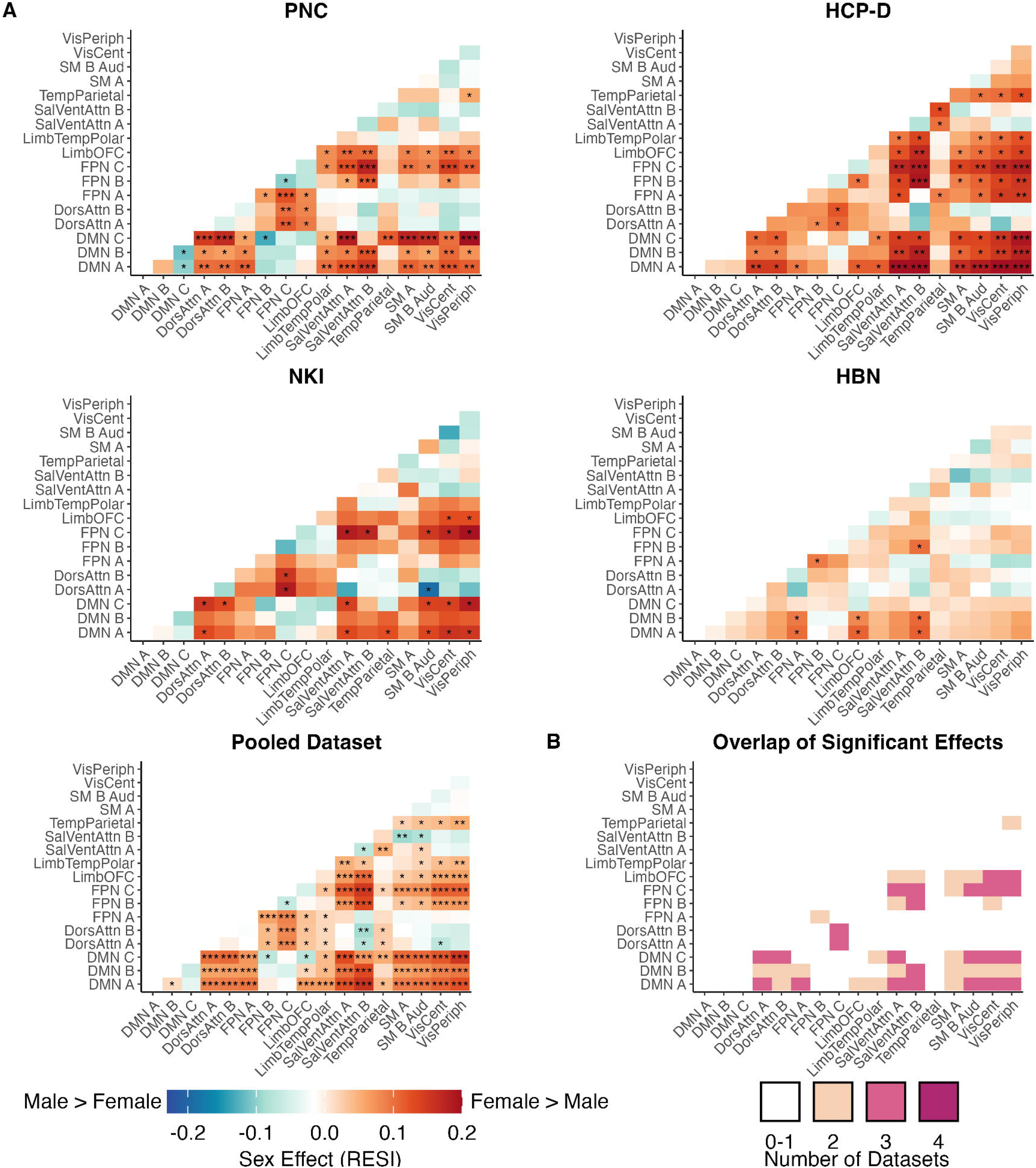
Females exhibit greater differentiation between higher- and lower-order networks along the principal gradient. **(A)** Between-network dispersion was quantified for each participant as the distance between pairs of network centroids along the principal functional connectivity gradient. Greater dispersion indicates greater network differentiation. Sex effects (RESI) of between-network dispersion for each pair of networks are shown for the PNC, NKI, HCP-D, HBN, and the pooled dataset. In every dataset, between-network dispersion is greater in females than in males for most network pairs (>70%). Across datasets, the largest effects are between higher-order association networks (default mode or frontoparietal control subnetworks) and lower-order salience/ventral attention and visual networks. Asterisks denote significance after FDR correction (\**Q*_FDR_ < 0.05, \*\**Q*_FDR_ < 0.001, \*\*\**Q*_FDR_ < 0.0001). **(B)** The matrix displays the number of datasets in which each network pair had a significant sex effect in between-network dispersion in either direction (*Q*_FDR_ < 0.05). Dispersion between default mode subnetwork A and salience/ventral attention subnetwork A, and between default mode subnetworks A and C and the lowest-order networks (somatomotor B/auditory, central visual, and peripheral visual), is significantly greater in females in every dataset except HBN. Network abbreviations as in Fig. 4.

## DISCUSSION

Using four independent, large-scale neuroimaging datasets, we delineated generalizable sex differences in functional brain organization in youth. Across datasets, we found that sex differences followed a consistent pattern along the sensorimotor-association (S-A) axis. Males showed greater connectivity between functional networks and between regions at opposite ends of the S-A axis. In contrast, females showed greater connectivity within networks, most prominently within default mode subnetwork A, and between regions at similar positions along the hierarchy. Beyond these connectivity differences, macroscale organization along the principal functional gradient also differed by sex, with females exhibiting a more expanded cortical hierarchy than males. Together, these findings reveal the cortical hierarchy as an overarching principle that unifies previously disparate observations of sex differences in human functional brain organization.

Opposing patterns of between- and within-network connectivity characterized regional- and network-level sex differences in functional connectivity, extending prior findings in youth and adulthood (*7*, *11*, *30*). The most reproducible effects converged on default mode subnetwork A, with females showing greater connectivity within this network and males showing greater connectivity between this network and lower-order attention and visual networks. Default mode subnetwork A includes medial prefrontal, posterior cingulate/precuneus, and inferior parietal cortex (*24*, *25*) and contributes to the core of the canonical default mode network (*31*, *32*). Sex differences involving default mode subnetworks B and C followed a similar pattern but were less consistent across datasets. Sex differences in default mode connectivity thus appear most robust in the network’s core. Functional independence of regions within the default mode network involved in self-referential processing and internally generated cognition from regions involved in perception and action has been proposed to support perceptually decoupled cognition (*33*). In this type of cognition, thought is less constrained by external sensory input and more guided by internal representations (*33*). Conversely, the default mode network may also integrate incoming external information with intrinsic memories and prior knowledge to construct context-dependent representations of ongoing experience (*34*). Our findings may therefore reflect sex differences in the degree to which the default mode network is segregated from versus integrated with externally oriented systems, although the cognitive consequences remain unknown. These findings identify higher-order association cortex as a key locus of sex differences in functional connectivity and point to a broader hierarchical organization of sex differences across the cortex.

Sex differences in functional connectivity were systematically organized along the cortical hierarchy. At the edge level, females showed greater connectivity within each end of the S-A axis, whereas males showed greater connectivity between regions spanning the axis. At the macroscale, females also showed greater dispersion between higher- and lower-order networks along the principal functional gradient, reflecting a more expanded cortical hierarchy. Because the principal functional gradient is derived from similarities in whole-brain connectivity profiles (*19*), the edge- and gradient-level findings are complementary. Greater connectivity within each end of the S-A axis and lower connectivity across the axis in females could render sensorimotor and association connectivity profiles more distinct than in males, consistent with greater separation along the principal gradient. This interpretation is consistent with recent large-scale lifespan work showing greater overall functional connectivity in males but greater functional system segregation in females, including within default mode and frontoparietal systems (*13*). Greater whole-brain and association-network segregation in females has also been reported in adulthood (*12*). Furthermore, our findings broadly align with Serio et al. (*21*), who identified sex differences in the S-A axis in young adults that were related to differences in connectivity profiles: connections within the default mode network were more prominent among the strongest connections in females, whereas somatomotor connections with other networks were more prominent among the strongest connections in males. However, they found no significant sex differences in between-network gradient dispersion. This discrepancy may reflect differences in age range as well as in gradient construction, including decomposition method, matrix thresholding, and network resolution. Together, our findings across regional-, network-, and edge-level connectivity as well as macroscale gradient organization consistently point to the cortical hierarchy as a common framework for sex differences in functional brain organization.

Sex differences in functional connectivity showed limited variation with age across youth, with age-by-sex interactions restricted to a small number of regions and network pairs. Because these interactions were detected only in the pooled analysis, age-related changes in sex differences are likely small and may require larger samples to detect reliably. Prior whole-brain studies in youth have similarly found few age-by-sex interactions, despite prominent sex differences or widespread developmental changes in functional connectivity (*7*, *35*). These findings raise the possibility that sex differences in functional organization emerge before the age range examined here, consistent with reports of sex-specific connectivity patterns as early as the first two years of life (*36*, *37*). Thus, the robust sex differences observed in our age window may partly reflect functional organization that is already differentiated earlier in development and subsequently follows relatively similar trajectories across childhood and adolescence. However, across a much broader age range, sex differences do appear to change: recent work spanning ages 8 to over 100 years found that sex differences in brain networks were smallest in late childhood and most pronounced in young to mid-adulthood (*30*). Although we observed few age-by-sex interactions in whole-brain connectivity, sex-specific developmental changes may still occur in specific circuits, such as amygdala connectivity during adolescence (*38*). Longitudinal studies spanning earlier and later developmental periods, ideally incorporating measures of pubertal maturation, will be needed to determine when these hierarchical sex differences emerge and how they change across the lifespan.

Several limitations of our work should be acknowledged. First, the cross-sectional design precludes inferences about within-person developmental trajectories, and our age range does not capture sex differences earlier in life. Second, we did not examine pubertal maturation or gonadal hormones, which may also contribute to sex differences in functional organization (*39–41*). We also did not investigate gender-related factors, which may have different associations than biological sex with functional brain organization (*16*). Third, the four datasets differed in sampling, acquisition, and participant characteristics, including enrichment for psychopathology in HBN. Although replication across heterogeneous datasets strengthens the generalizability of our findings, these differences may also contribute to variation in effect magnitude across datasets. Notably, sex differences in macroscale gradient organization were weaker in HBN. Whether this reflects the clinical characteristics of this sample or other dataset-specific factors remains to be determined. Fourth, our analyses used a group-level functional atlas, which does not capture individual variation in network topography, a feature that differs by sex, particularly in association cortex (*5*, *10*). Because between- and within-network connectivity depend on how regions are assigned to networks, individualized functional networks may provide a more precise characterization of these effects. Finally, the cognitive and clinical significance of these sex differences remains undetermined, as the present study characterizes differences in functional organization without establishing their consequences for behavior or risk for psychopathology.

In conclusion, across four independent developmental datasets, sex differences in functional brain organization were consistently organized along the cortical hierarchy. Females exhibited greater segregation and expansion of the hierarchy, whereas males showed greater integration across the S-A axis and a more compressed hierarchy. By identifying the cortical hierarchy as a common framework for these differences, this work moves beyond cataloging localized sex differences to reveal their broader organization across the cortex. Moving forward, characterizing reproducible variation in hierarchical brain organization across development may provide a foundation for understanding individual differences in cognition and neuropsychiatric risk across the lifespan.

## MATERIALS AND METHODS

### Participants

We analyzed resting-state and task fMRI data from four large-scale developmental datasets (total *N* = 3,355; 1,608 female): the Philadelphia Neurodevelopmental Cohort (PNC; *n* = 1,207; 646 female; ages 8–23), the Nathan Kline Institute-Rockland Sample (NKI; *n* = 397; 186 female; ages 6–22), the Human Connectome Project: Development (HCP-D; *n* = 625; 337 female; ages 5–22), and the Healthy Brain Network (HBN; *n* = 1,126; 439 female; ages 5–22). These samples are identical to those in our prior work on the development of functional connectivity along the S-A axis (*20*). Inclusion and exclusion criteria are summarized briefly below, with full details and dataset demographic characteristics reported previously (see Table 1 of (*20*)).

The PNC (*42*, *43*) is a community sample of children, adolescents and young adults from the greater Philadelphia area recruited to study brain development. Study procedures were approved by the Institutional Review Boards of the University of Pennsylvania and the Children’s Hospital of Philadelphia. NKI (*44*, *45*) is a community-ascertained lifespan sample (ages 6–85) designed to reflect the demographic composition of the United States; we included participants aged 6–22. NKI procedures were approved by the Institutional Review Board of the Nathan Kline Institute. HCP-D (*46*) recruited typically developing children and adolescents across four sites (University of Minnesota, Harvard University, Washington University in St. Louis, and University of California, Los Angeles), with a sampling design that reflects the demographics of youth in the United States. HCP-D procedures were approved by a central Institutional Review Board at Washington University in St. Louis. HBN (*47*) is a community-referred sample of children and adolescents from the New York City area. Families with concerns about their child’s psychiatric symptoms were encouraged to participate. HBN procedures were approved by the Chesapeake Institutional Review Board (now Advarra, Inc.). In all datasets, written informed consent was obtained from participants aged 18 or older. For participants under 18, a parent or legal guardian provided written informed consent and the participant provided assent.

Sex was recorded as female or male in the PNC, NKI, HCP-D, and HBN. In the PNC, demographic information was caregiver-reported for participants younger than 18 years and self-reported for those aged 18 years and over (*43*). In NKI, sex was caregiver-reported for participants aged 6-11 years and self-reported for those aged 12 years and older (*44*). In HCP-D, sex assigned at birth was parent-reported and self-reported for those aged 18 years and older (*46*). In HBN, sex assigned at birth was recorded, with demographic information caregiver-reported for participants aged 5-17 years and self-reported for those aged 18 years and older (*47*). HCP-D additionally assessed self-perceived gender, including nonbinary options. Comparable gender identity measures were not documented in the other datasets.

### Sample construction

The initial samples included 1,559 participants with fMRI data in the PNC, 1,288 in NKI, 652 in HCP-D, and 2,255 in HBN. Exclusion and quality assurance criteria were applied in the following order. First, we included participants aged 5-23 years for each dataset. We excluded 862 participants from the NKI lifespan sample; no participants were excluded on age in the PNC, HCP-D, or HBN. Second, we excluded participants with medical conditions that could affect brain function (e.g., malignancy, systemic illness, neurological disorders, and brain injury) or with gross neurological abnormalities, where these data were available (PNC, *n* = 146; HCP-D, *n* = 21). Third, we excluded participants whose T1-weighted images failed manual quality assurance. In the PNC, three trained raters assessed image quality (*n* = 39 excluded). In NKI and HBN, raters used the Swipes for Science web application (*48*) to pass or fail images on overall quality and gray-white matter boundary definition (NKI, *n* = 5; HBN, *n* = 586 excluded). T1-weighted quality assurance for HCP-D was performed by the data-collection team.

Fourth, as in prior work, we excluded individual fMRI runs with mean framewise displacement (FD) ≥ 0.3 mm. Participants were excluded if all fMRI scans failed head motion exclusion (PNC, *n* = 112; NKI, *n* = 18; HCP-D, *n* = 2; HBN, *n* = 354). In the PNC, HCP-D, and HBN, retained resting-state and task runs were concatenated to maximize scan time. Because NKI acquired only resting-state fMRI, only these scans were used. For NKI participants with multiple imaging sessions, we used the session with the most runs that passed T1 and head motion quality control. Finally, we excluded participants with less than 7 minutes of retained fMRI data (PNC, *n* = 55; NKI, *n* = 6; HCP-D, *n* = 4; HBN, *n* = 189). A flow chart of sample construction for each dataset is provided in our prior report using this sample (see Supplementary Figure 16 of (*20*)).

### Image acquisition and processing

Acquisition parameters and image processing for T1-weighted and functional MRI in each dataset have been previously reported (*20*) and are summarized here. Minimal preprocessing used fMRIPrep (*49*) version 20.2.3 (PNC and NKI) or 22.0.2 (HCP-D and HBN). The newer release was used for HCP-D and HBN to support susceptibility distortion correction with reverse phase-encoding field maps.

T1-weighted images were corrected for intensity non-uniformity with ANTs (2.3.3) N4BiasFieldCorrection (*50*, *51*), skull-stripped with the ANTs brain extraction workflow, and segmented into tissue classes with FSL FAST 5.0.9 (PNC and NKI) and 6.0.5.1 (HCP-D and HBN) (*52*). Cortical surfaces were reconstructed with FreeSurfer 6.0.1 (PNC and NKI) or 7.2.0 (HCP-D and HBN) (*53*). Each T1-weighted image was normalized to MNI152NLin6Asym and MNI152NLin2009cAsym space by nonlinear registration with ANTs.

For functional data, a B0 field map was estimated from a dual-echo gradient-recalled echo phase-difference map (PNC) or from reverse phase-encoding EPI references with FSL TOPUP (HCP-D and HBN). Susceptibility distortion correction was not performed for NKI, which lacked field maps, or for the 34 PNC participants (72 runs) and 6 HBN participants (13 runs) without field maps. The BOLD reference was co-registered to the T1-weighted image with boundary-based registration (FreeSurfer bbregister, six degrees of freedom). Head-motion parameters were estimated with FSL mcflirt (*54*), and slice-timing correction was performed with AFNI 3dTshift (*55*). Motion correction, distortion correction and spatial transforms were applied in a single resampling step. BOLD time series were then projected to the fsLR 32k surface as 91k-grayordinate files (*56*).

Postprocessing used XCP-D (*57–59*) version 0.0.8 (NKI), 0.3.0 (HCP-D), or 0.3.2 (PNC and HBN). Framewise displacement was computed with a 50 mm head radius (*60*). BOLD data were despiked, mean-centered, and linearly detrended. We then regressed 36 nuisance parameters from the data: six motion parameters, mean global signal, mean white matter signal, mean cerebrospinal fluid signal, their temporal derivatives, and quadratic expansions of these terms (*58*, *61*). Denoised time series were parcellated with the Schaefer 200-parcel atlas (*24*). Retained resting-state and task time series were concatenated for each participant, and the Pearson correlation between the concatenated time series of every pair of cortical regions yielded a 200-by-200 functional connectivity matrix. This concatenation approach is supported by evidence that functional networks are largely shared across task and rest states, that individual rather than task-related variability accounts for most variation in functional connectivity (*62*), and that longer scan time improves the reliability of connectivity estimates (*63*, *64*). In-scanner head motion was quantified for each participant as mean FD averaged across all retained runs.

### Cortical parcellation, functional networks, and the sensorimotor-association axis

We used the Schaefer 200-parcel atlas (*24*), with each region assigned to one of 17 functional networks from the Yeo 17-network solution (*25*). The networks included the central visual, peripheral visual, somatomotor A, somatomotor B/auditory, dorsal attention A and B, salience/ventral attention A and B, limbic temporal pole, limbic orbitofrontal, frontoparietal control A, B, and C, default mode A, B, and C, and temporal parietal.

To place regions within the cortical hierarchy, we used the sensorimotor-association (S-A) axis (*17*). The S-A axis was derived by rank-ordering vertices on each of ten cortical feature maps that vary systematically between primary sensorimotor and transmodal association cortex, then averaging the ranks across maps. The vertex-level axis (https://github.com/PennLINC/S-A_ArchetypalAxis) was parcellated with the Schaefer 200 atlas, giving each region a unique rank from 1 (most sensorimotor) to 200 (most associative).

### Quantification of functional connectivity measures

To examine sex differences in functional connectivity, we quantified region-, network-, and edge-level functional connectivity. All measures used signed, unthresholded Pearson correlations. At the region level, functional connectivity (FC) strength was defined as the mean correlation between a region and all other 199 cortical regions. To understand which connections drove sex differences in FC strength, we decomposed this measure into two components.

Between-network connectivity was computed as the mean correlation between a region and all regions outside its functional network. Within-network connectivity was quantified as the mean correlation between a region and all other regions in its own network. At the network level, we computed connectivity for each pair of the 17 functional networks. Connectivity between two networks was the average correlation between every region in one network and every region in the other. Connectivity within a network was the average correlation among all regions in that network. This yielded 153 total network pairs: 136 pairs of distinct networks and 17 within-network pairs. At the edge level, we analyzed the connectivity between each of the 19,900 unique pairs of regions.

### Harmonization of multi-site data

To ensure imaging measures were comparable across sites, connectivity measures in multi-site datasets (HCP-D and HBN) were harmonized across sites. We used an extension of Correcting Covariance Batch Effects (CovBat) (*65*) that models age as a smooth term in a generalized additive model (GAM) during both the mean-correction and covariance-correction stages, similar to ComBat-GAM (*66*). Age was modeled with an unpenalized thin plate regression spline (*k* = 3), matching the models described below. Sex and in-scanner motion were included as linear covariates so that variation associated with them was preserved during harmonization. Harmonization was implemented with the *ComBatFamily* package (version 0.2.1) in R 4.1.2. We chose CovBat because functional connectivity data show covariance-related batch effects (*65*, *67*), and because the GAM formulation respects the non-linear age effects in our models.

We harmonized each region-level measure (FC strength, between-network connectivity, and within-network connectivity), network-pair connectivity, and edge-level connectivity. For the gradient analyses, we also harmonized each participant’s full connectivity matrix. To do this, we vectorized the upper triangle (19,900 edges), harmonized the edges together, and rebuilt symmetric 200-by-200 matrices.

To increase statistical power, we also analyzed a pooled dataset that combined all participants from the four datasets (*N* = 3,355). In the pooled dataset, each combination of dataset and acquisition site was treated as a separate batch: PNC, NKI, each HCP-D site, and each HBN site. This harmonized measures across both datasets and sites in one step, using the same CovBat-GAM approach above.

### Statistical models of sex differences

To estimate sex differences while accounting for developmental change, we fit GAMs with the *mgcv* package (version 1.8.39) in R (*68*, *69*). Separate models were fit for each region, network pair, and edge, in each dataset and in the pooled dataset. The connectivity measure was the dependent variable, age was a smooth term, and sex and in-scanner motion (mean FD) were linear covariates. Specifically, age was included as a smooth term to model linear and nonlinear changes in functional connectivity across the age range of the sample, allowing sex differences to be estimated after accounting for age-related variation in connectivity.

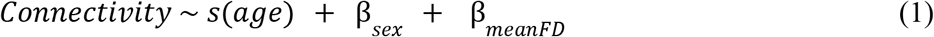

As in our prior work (*20*), age was modeled with an unpenalized thin plate regression spline with a maximum basis complexity (*k*) of 3 to avoid overfitting. Sex was coded with males as the reference level. Positive effect sizes indicate greater values in females and negative effect sizes indicate greater values in males. Significance of the sex effect was assessed with the *p*-value for the parametric sex coefficient from each GAM (a two-sided Wald *t*-test, as implemented in *mgcv*). *P*-values were corrected for multiple comparisons using the false discovery rate across all regions (*n* = 200), network pairs (*n* = 153), or edges (*n* = 19,900), separately for each measure and dataset (*Q*_FDR_ < 0.05). To test whether sex differences varied with age, we added an age-by-sex factor-smooth interaction to Eq. 1:

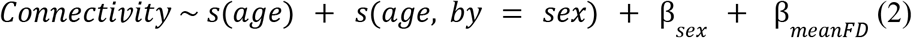

Sex was entered as an ordered factor with males as the reference level, so *s*(age, by = sex) estimates the difference between the female and male age smooths, with the same basis settings as above. We assessed the interaction between age and sex with an F-test of this term and applied FDR correction as above. Interactions were examined in each dataset and in the pooled dataset. To visualize developmental trajectories, we generated predicted values from Eq. 2 separately for females and males at 200 evenly spaced ages across each dataset’s age range, with motion held at its median.

### Effect size estimation

We quantified sex effects with the robust effect size index (RESI) (*22*, *23*). RESI is a standardized effect size that can be computed for any model parameter, is comparable across model types, and does not depend on sample size. For reference, under equal group sizes and equal within-group variances, RESI is equivalent to half of Cohen’s *d* (e.g., RESI = 0.1 corresponds to *d* = 0.2). Following prior work (*23*), we interpreted absolute RESI values of 0-0.1, 0.1-0.25, 0.25-0.4, and > 0.4 as no-to-small, small-to-medium, medium-to-large, and large effects, respectively. These ranges correspond to conventional benchmarks for Cohen’s *d* (*70*).

RESI was computed from the *t*-statistic for the sex coefficient in Eq. 1 using the *t2S* function in the *RESI* package (version 1.3.2; (*23*)) in R. For region-level, network-level, and gradient dispersion measures, this *t*-statistic was based on a robust (HC0) sandwich covariance estimate, which does not assume equal residual variance across participants. For edge-level connectivity, RESI was computed from the model-based *t*-statistic from each GAM to reduce computation across 19,900 edges. Statistical significance was assessed separately using p-values from the original GAMs.

In a post hoc analysis, we assessed whether replicated effects were also the largest. We ranked regions and network pairs by mean RESI across datasets, separately for male-greater and female-greater effects, to examine whether those significant in all four datasets fell within the top 20%.

### Spin-based spatial null models

To account for spatial autocorrelation in cortical maps, we assessed correspondence between maps with spin-based spatial null models (“spin tests”) (*28*). Spin tests generate a null distribution by rotating spherical projections of a cortical map, which preserves its spatial covariance structure. We generated 10,000 rotations of Schaefer 200 region centroids on the fsaverage sphere with the “rotate_parcellation” algorithm (*71*) and used the same rotations for all spin tests. For correlations between two regional maps, *p*_spin_ was the proportion of null correlations exceeding the observed correlation (*71*).

### Alignment of edge-level sex effects with the S-A axis

To test whether sex differences in edge-level connectivity were organized along the cortical hierarchy, we labeled each edge with the S-A axis ranks of the two regions it connects. We then modeled the edge-wise sex effect (RESI) as a function of these two ranks using a tensor product smooth (*k* = 3) in *mgcv* and visualized the fitted surface as a topographical plot (*72*, *73*):

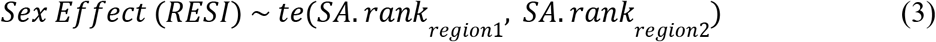

Because the ordering of an edge’s two regions is arbitrary, each edge was included with its regions in both orders, making the fitted surface symmetric. To test whether S-A ranks explained significant variance in edge-level sex effects, we used the adjusted *R*² of Eq. 3 as the test statistic. We rotated the S-A axis 10,000 times, reassigned the rotated ranks to each edge’s regions, and refit Eq. 3 to generate a null distribution of adjusted *R*² values. Here, *p*_spin_ was the proportion of rotations whose adjusted *R*² equaled or exceeded the observed adjusted *R*².

### Meta-analytic decoding with Neurosynth

To relate regions showing replicable sex differences to cognitive functions, we compared regional maps of sex differences with meta-analytic activation maps from Neurosynth (*26*). Following prior work (*74*, *75*), we selected 125 terms from the Neurosynth database (version 0.7) that correspond to cognitive concepts in the Cognitive Atlas (*27*). The full list of cognitive terms is provided in prior work (see Supplementary Table 2 of (*75*)). For each term, we used *NiMARE* (version 0.21.0, (*76*)) in Python (version 3.11) to generate an association *z*-statistic map with a multilevel kernel density chi-square analysis, using default parameters (*77*). Values in these maps indicate how consistently a region was activated in studies mentioning the term relative to studies that did not. Maps were parcellated into the Schaefer 200 atlas with *neuromaps* (version 0.0.7) (*78*) and *z*-scored across regions.

For between- and within-network connectivity, we built a sex-effect map from regions with an FDR-significant sex difference in two or more datasets, using each region’s RESI averaged across the PNC, NKI, HCP-D, and HBN. We computed Spearman’s rank correlations between each sex-effect map and each term map, assessed significance with spin tests, and applied FDR correction across the 125 terms. We focused on significant terms in the direction of interest for each measure: negative correlations for between-network connectivity (greater in males) and positive correlations for within-network connectivity (greater in females). These correlations indicate spatial correspondence with task-activation maps; importantly, they should *not* be interpreted to imply sex differences in cognitive function or performance in these domains.

### Principal gradient of functional connectivity and dispersion analysis

To test whether the cortical hierarchy was expanded in females and compressed in males, we computed the principal gradient of functional connectivity (*19*) for each participant with *BrainSpace* (version 0.2.1) (*79*) in Python. Gradients were computed in each dataset and in the pooled dataset. Harmonized connectivity matrices were used for HCP-D, HBN, and the pooled dataset.

First, we built a group template for each dataset by averaging participants’ connectivity matrices. We converted the group matrix to an affinity matrix with the normalized angle kernel, without thresholding, and decomposed it with principal component analysis (PCA), retaining 10 components. Because the sign of a principal component is arbitrary, we oriented the first principal component (PC1) to correlate positively with the S-A axis, so that higher loadings indicate positions closer to the association end of the hierarchy. Next, we computed gradients for each participant in the same way and aligned them to the group template with a single iteration of Procrustes rotation across all 10 components (*79*), which placed all participants’ gradients in a common space. In all datasets and in both sexes, the aligned principal gradient spanned sensorimotor to association cortex (**Fig. S5**).

To characterize how expanded or compressed each participant’s principal gradient was, we then quantified the spread of regions along their aligned PC1 (*21*, *29*). For a given set of regions, we defined its centroid as the median PC1 loading of those regions (i.e., their center along the gradient). Global dispersion captured how widely all regions were spread around the center of the gradient and was computed as the sum of squared distances between each region’s PC1 loading and the centroid of all 200 regions. Greater global dispersion indicates a more expanded gradient. Each network’s centroid was similarly defined as the median PC1 loading of that network’s regions. Between-network dispersion for each of the 136 pairs of networks was the absolute difference between the two networks’ centroids along PC1. Greater between-network dispersion indicates greater differentiation between networks along the principal gradient.

We modeled sex differences in global and between-network dispersion with the GAM in Eq. 1 and computed the RESI sex effects as described above. As with the connectivity measures described previously, significance of the sex effect was assessed with the *p*-value for the parametric sex coefficient from each GAM (a two-sided Wald *t*-test, as implemented in *mgcv*). For between-network dispersion, *p*-values were corrected for multiple comparisons using the false discovery rate across the 136 network pairs, separately for each dataset (*Q*_FDR_ < 0.05).

## Author contributions

Conceptualization: A.C.L., S.S., T.D.S. Methodology: A.C.L., M.D.H., S.N.V., S.S., T.D.S. Software: A.C.L. Validation: M.D.H. Formal analysis: A.C.L. Investigation: A.C.L. Resources: A.C.L., M.C., A.R., D.M.B., A.R.F., R.E.G., R.C.G., G.K., M.P.M., L.H.S., T.D.S. Data curation: A.C.L., M.C., T.S. Visualization: A.C.L. Supervision: S.S., T.D.S. Writing—original draft: A.C.L. Writing—review and editing: A.C.L., M.D.H., A.F.A.-B., J.B., D.M.B., M.C., E.A.F., A.R.F., R.E.G., R.C.G., A.S.K., G.K., M.P.M., L.P., A.R., T.S., G.A.S., G.S., R.T.S., L.H.S., V.J.S., S.N.V., X.Z., S.S., T.D.S.

## Use of artificial intelligence

ChatGPT (OpenAI) was used to assist with manuscript revision to improve the clarity, readability, and flow of the text. All resulting text was subsequently reviewed and edited by the authors, who take full responsibility for the content of the manuscript.

## Data and code availability

This study analyzed publicly available data. The Philadelphia Neurodevelopmental Cohort is available from the Database of Genotypes and Phenotypes (phs000607.v3.p2; https://www.ncbi.nlm.nih.gov/projects/gap/cgi-bin/study.cgi?study_id=phs000607.v3.p2). The Nathan Kline Institute-Rockland Sample is available from OpenNeuro (https://openneuro.org/datasets/ds001021/versions/1.0.0). The Human Connectome Project: Development is available through the NIMH Data Archive (https://nda.nih.gov/). The Healthy Brain Network is available at https://fcon1000.projects.nitrc.org/indi/cmihealthybrainnetwork/.

All analysis code is available at https://github.com/PennLINC/luo_sex_diff. A detailed guide to the code and procedures for replicating all analyses is available at https://pennlinc.github.io/luo_sex_diff. R analyses were run in a publicly available container (docker://audreycluo/r-packages-for-parcc:0.0.1), and the Python environment specification is provided in the repository. Figures were generated in R (version 4.4.3) with *ggplot2* (version 3.5.2), *ggseg* (version 1.6.5), and *ggsegSchaefer* (version 2.0.4).

