## Supplementary Information for "Generalizable sex differences in functional brain organization follow the cortical hierarchy in youth"

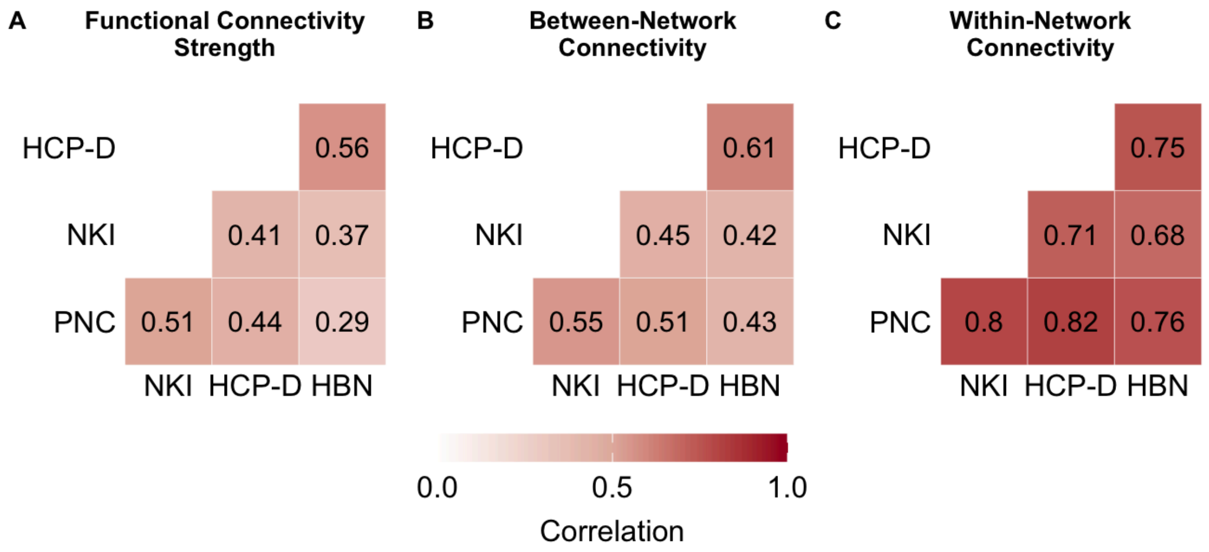


**Fig. S1. Sex effects on region-level connectivity are spatially correlated across datasets.** Pearson correlations between regional sex-effect maps (RESI) for each pair of datasets are shown for **(A)** functional connectivity strength, **(B)** between-network connectivity, and **(C)** within-network connectivity. All correlations were significant as assessed by spin-based spatial permutation tests (all $p_{spin} < 0.001$).

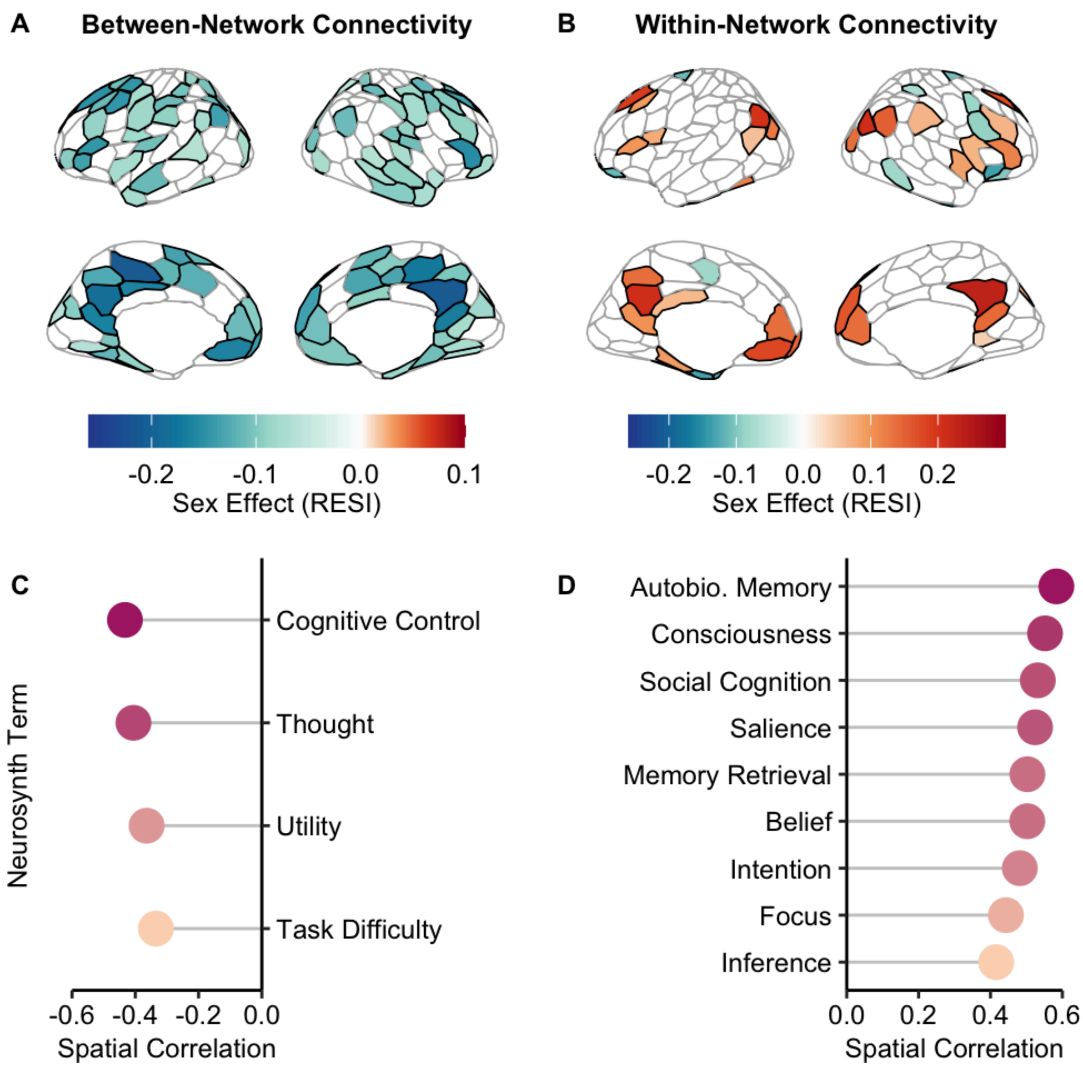


**Fig. S2. Sex differences in between- and within-network connectivity spatially align with meta-analytic maps of cognitive terms. (A and B)** Regions with a significant sex difference ($Q_{FDR} <$ 0.05) in two or more datasets are shown for between-network connectivity **(A)** and within-network connectivity **(B)**, colored by the sex effect (RESI) averaged across the PNC, NKI, HCP-D, and HBN. Between-network sex effects are widespread and reflect greater connectivity in males (blue; negative RESI), whereas within-network effects predominantly reflect greater connectivity in females (red; positive RESI). **(C)** Neurosynth terms significantly correlated with the between-network sex effect ($Q_{FDR}$ < 0.05): cognitive control, thought, utility, and task difficulty. These terms are most associated with cortical regions where males display greater between-network connectivity ($r$ = -0.43 to -0.33). **(D)** Neurosynth terms significantly correlated with the within-network sex effect, including autobiographical memory, social cognition, and belief. These terms are most associated with cortical regions where females show greater within-network connectivity ($r$ = 0.42 to 0.58). **(C)** and **(D)** show all terms surviving FDR correction. Spearman's rank correlations were used to quantify the association between sex effects in the regions shown in **(A)** and **(B)** and the meta-analytic map for each of 125 Neurosynth terms selected using the Cognitive Atlas. Statistical significance was assessed using spin tests and FDR correction across terms ($Q_{FDR}$ < 0.05). These correlations index spatial correspondence with task-activation maps and do not imply sex differences in the corresponding cognitive functions or performance. Autobiographical is abbreviated as autobio.

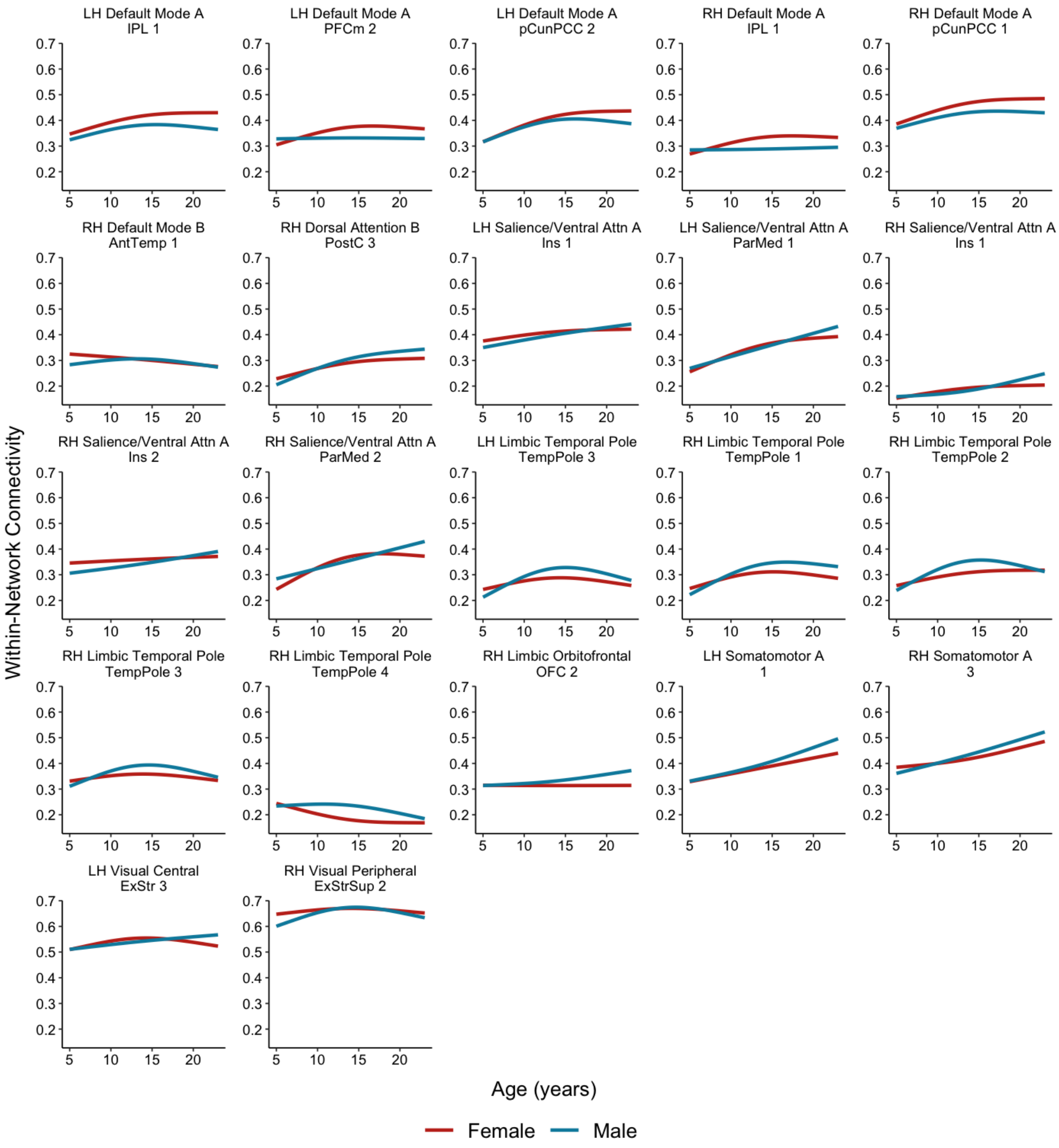


**Fig. S3. Age-by-sex interactions in within-network connectivity.** Developmental trajectories of within-network connectivity for females (red) and males (blue) are shown for the 22 regions with a significant age-by-sex interaction in the pooled dataset ($Q_{\mathrm{FDR}} < 0.05$). Interactions were evaluated with generalized additive models fit to harmonized data that included a smooth term for age, an age-by-sex factor-smooth interaction, and sex and in-scanner motion as covariates. Lines show model-predicted values from ages 5 to 23, with in-scanner motion held at its median. No region showed a significant age-by-sex interaction for FC strength or between-network connectivity in the pooled dataset. LH, left hemisphere; RH, right hemisphere; Attn, attention; ExStr, extrastriate; ExStrSup, superior extrastriate; IPL, inferior parietal lobule; ParMed, medial parietal; PFCm, medial prefrontal cortex; pCunPCC, precuneus/posterior cingulate cortex; AntTemp, anterior temporal cortex; PostC, postcentral cortex; Ins, insula; TempPole, temporal pole; OFC, orbitofrontal cortex.

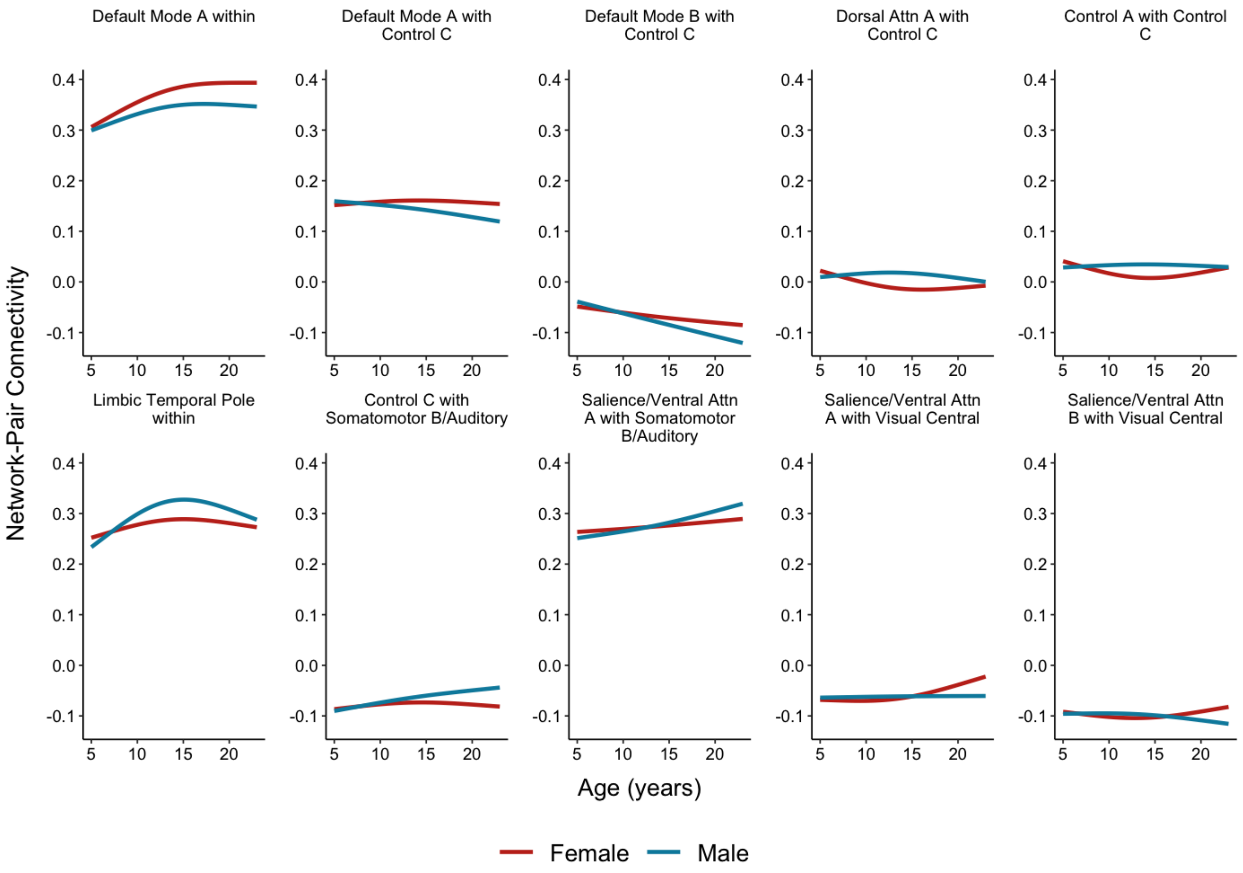


**Fig. S4. Age-by-sex interactions in network-pair connectivity.** Developmental trajectories of network-pair connectivity for females (red) and males (blue) are shown for the 10 network pairs with a significant age-by-sex interaction in the pooled dataset ($Q_{FDR} < 0.05$). Models, predicted values, and abbreviations are as in **Fig. S3**.

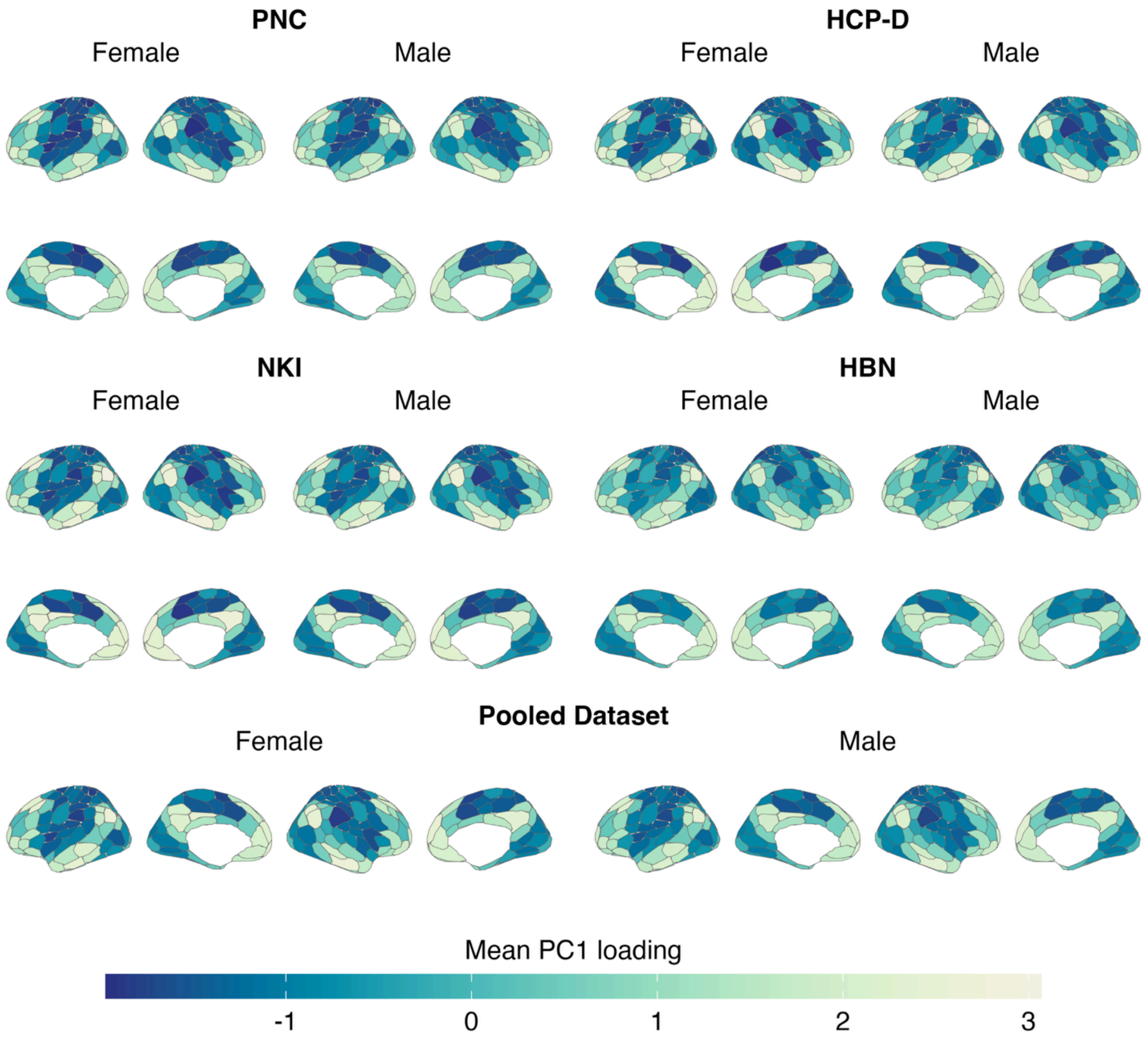


**Fig. S5. The principal gradient of functional connectivity in females and males.** Mean regional loadings on the first principal component (PC1) of functional connectivity are shown separately for females and males in the PNC, NKI, HCP-D, HBN, and the pooled dataset. In all datasets and in both sexes, the principal gradient spans from sensorimotor cortices (negative loadings) to association cortices (positive loadings).

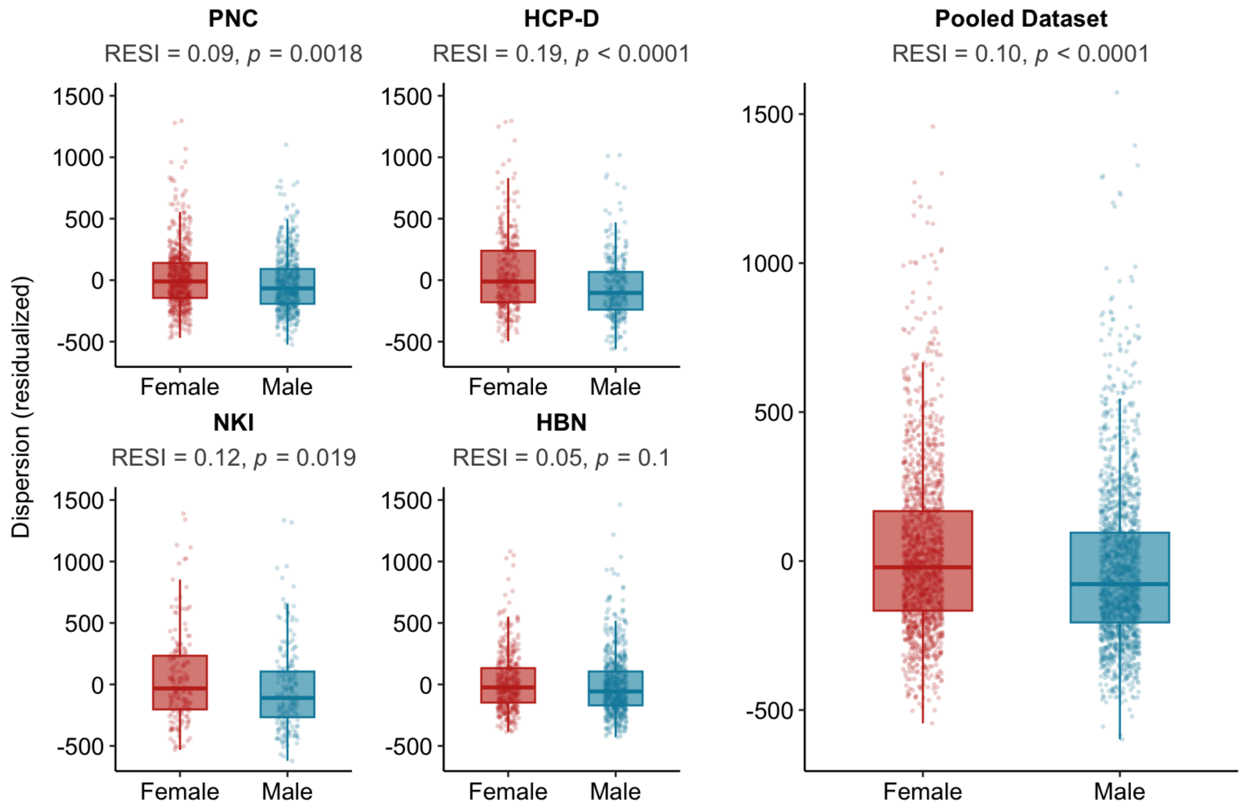


**Fig. S6. Global dispersion along the principal gradient is greater in females.** Global dispersion, defined as the overall spread of regions along the principal functional connectivity gradient, is shown for females and males in the PNC, NKI, HCP-D, HBN, and the pooled dataset. Values are residualized for age and in-scanner motion. Sex effects (RESI) and *p*-values from generalized additive models are shown above each panel. Global dispersion is greater in females in all datasets except HBN. Boxes indicate the median and interquartile range; points represent individual participants.
